## Supplementary Information for "Insight into the structural basis for dual nucleic acid - recognition by the Scaffold Attachment Factor B2 protein"

**This document contains:**

- 1 Supplementary Table
- 12 Supplementary Figures

Supplementary Table 1: Overview of DNA oligos of this study used to generate SAFB2 SAP domain variants.

| Name | Sequence 5‘ → 3‘ | Usage |
|---|---|---|
| SAP_R45A_fw | CCGTGCGGAACTTAAAAAGGCGAATCTGGATACCGGTGG | SPRINP[(a)] primers to generate single-point mutant $SAP_{R45A}$ |
| SAP_R45A_rev | CCACCGGTATCCAGATTCGCCTTTTTAAGTTCCGCACGG | |
| SAP_K53A/S54A_fw | CGCAATCTGGATACCGGTGGTAACGCAGTTCTGATGGAGCGCCTG | SPRINP[(a)] primers to generate double-point mutant $SAP_{K53A/S54A}$ |
| SAP_K53A/S54A_rev | CAGGCGCTCCATCAGAACTGCTGCGTTACCGGTATCCAGATTGCG | |
| SAP_S31A_fw | GACTCGTCGGTTAGCGGAGTTGCGCG | SDM[(b)] primers to generate single-point mutant $SAP_{S31A}$ |
| SAP_S31A_rev | CCCGTTTCGGCCACGCCC | |
| SAP_T26_fw | ACTCGTCGGTTATCGGAGTTGCGC | SDM[(b)] primers to generate N-terminal shortened $SAP_{26-70}$ |
| SAP_T26_rev | GCCCTGAAAATAAAGATTCTCAGAACCACTGCC | |

(a) Site-directed mutagenesis by SPRINP method, according to Edelheit et al; 2009 [1]. SPRINP, Single-Primer Reactions IN Parallel
(b) Cloning according to manufacturer’s protocol included in NEBaseChanger®, New England Biolabs. SDM, site-directed mutagenesis

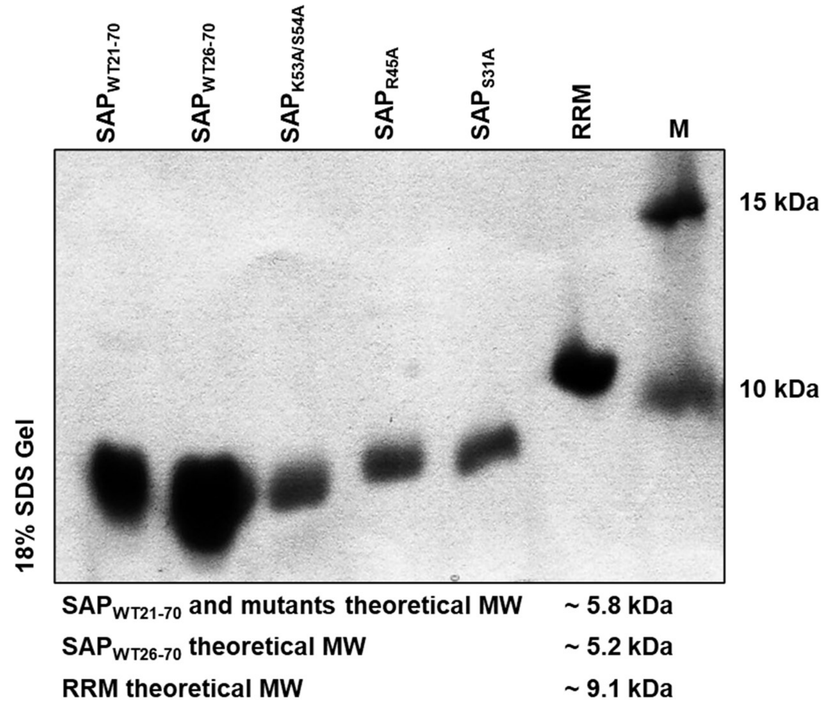


**Supplementary Figure 1:** SDS-gel (18% acrylamide) showing representative aliquots of all proteins used in this study ($SAP_{WT21-70}$, $SAP_{WT26-70}$, $SAP_{K53A/S54A}$, $SAP_{R45A}$, $SAP_{S31A}$, and RRM) from final purified samples.

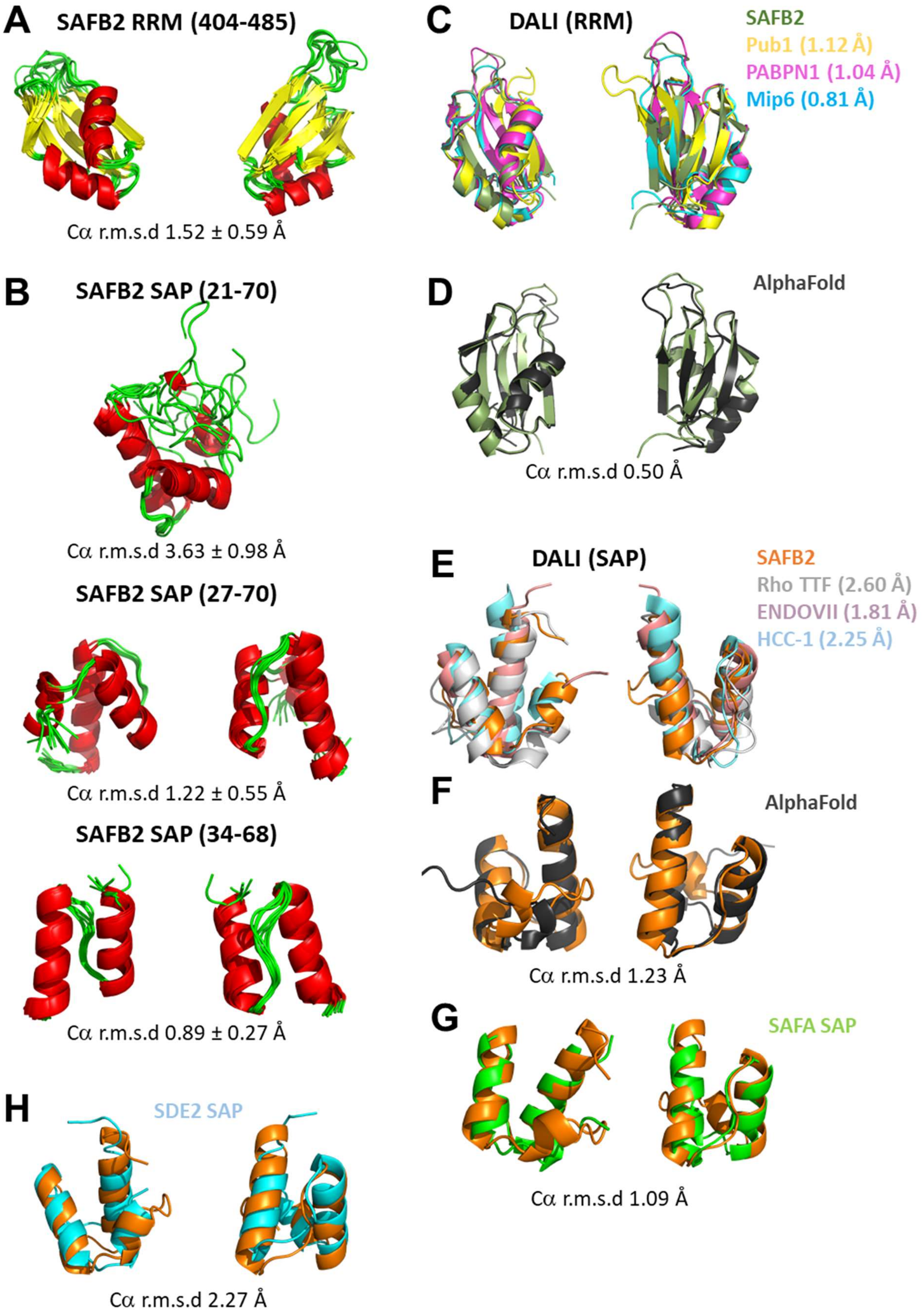


**Supplementary Figure 2:** CS-Rosetta derived structural models. **(A)** Ensemble of best 10 structures obtained from CS-Rosetta [2] for the SAFB2 RRM. The RMSD value to the lowest energy structure is given as average and standard deviation. **(B)** The same is in panel A for the SAFB2 SAP domain. The ensemble is shown as structural alignment for the full NMR construct (top), the N-terminally truncated version (middle, see also Suppl. Fig. 3) and two-helical core domain (bottom), indicating increasing convergence. **(C, D)** Comparison of the CS-Rosetta-derived lowest energy SAFB2 RRM domain structure with the three best hits obtained from a DALI [3] search (C) or with an AlphaFold [4] derived model of the identical boundaries (D). Individual RMSD values are given. Underlying PDB codes are: 3MD3 (Pub1, unpublished), 3B4M (PABPN1 [5]), 5D77 (Mip6, unpublished). **(E-H)** Comparison of the CS-Rosetta-derived lowest energy SAFB2 SAP domain structure with either the three best hits obtained from a DALI [3] search (E), with an AlphaFold derived model of the identical boundaries (F), with the SAP domain of SAFA, or with the SAP domain of ssDNA-binding SDE2 using analogous boundaries (G). Individual RMSD values are given. Underlying PDB codes are: 5JJI (Rho TTF [6]), 1E7D (ENDOVII [7]), 7N99 (SDE2) [8], 2DO1 (HCC-1, unpublished).

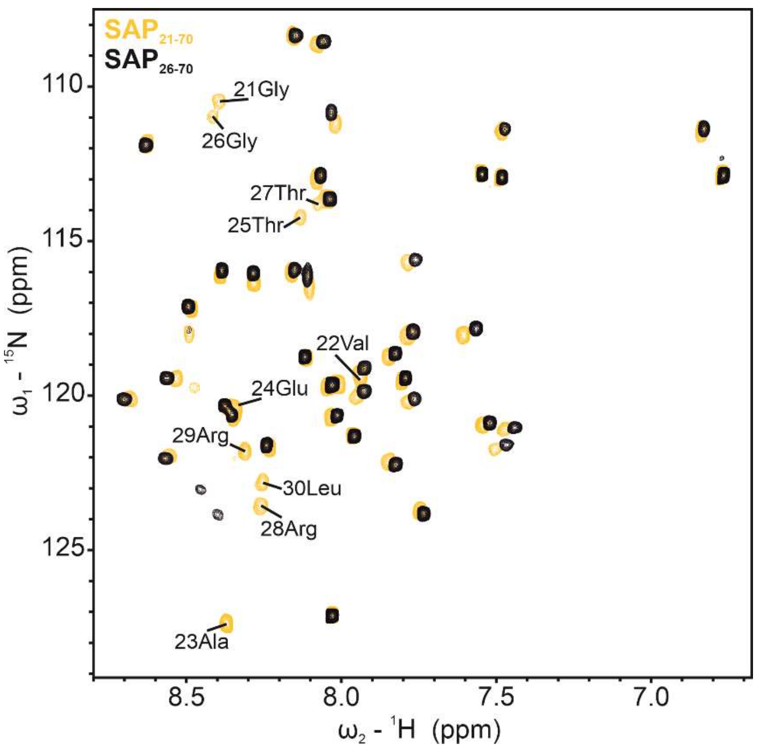


**Supplementary Figure 3:** N-terminal truncation of SAFB2 SAP does not affect the domain's core fold. Overlay of $^{1}H$-$^{15}N$-HSQC spectra of wild-type $SAP_{21-70}$ (yellow) and $SAP_{26-70}$ (black). N-terminal residues are labelled with their assignments for orientation. The new Gly26 amide of $SAP_{26-70}$ is line-broadened beyond detection. Only residues of the N-terminal extension within the truncated and non-truncated SAP versions are affected with respect to each other as seen by their perturbed chemical shifts.

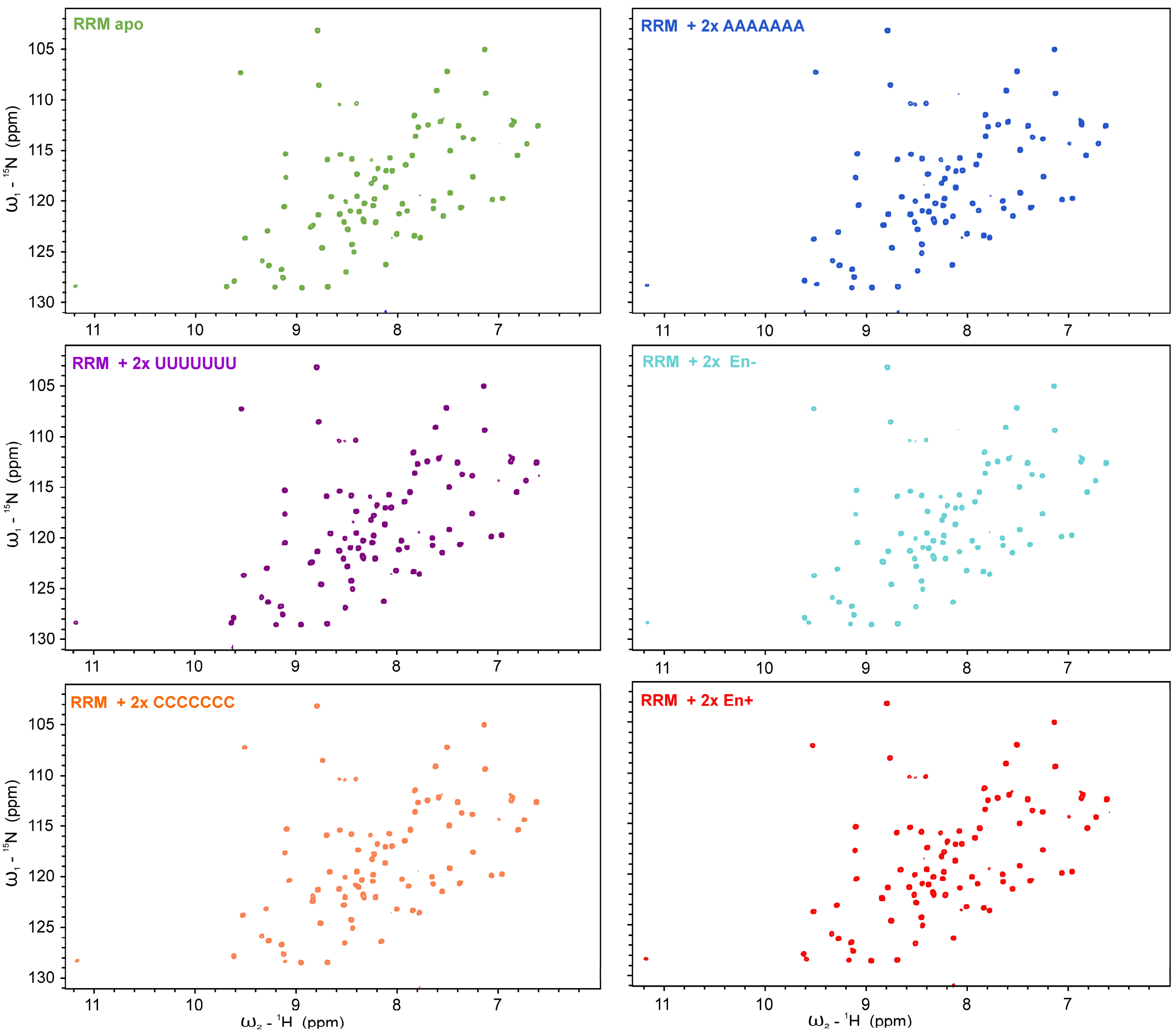


**Supplementary Figure 4:** RNA preferences of the SAFB2 RRM domain. Full $^{1}H$-$^{15}N$ HSQC spectra of the apo RRM (green, top left) and the RRM after addition of 2x molar excess of either $U_7$ (purple, middle left), $C_7$ (orange, bottom left), $A_7$ (blue, top right), En+ (cyan, middle right) or En- (red, bottom right). Spectra correspond to zoom-ins in main text Figure 4.

**A**

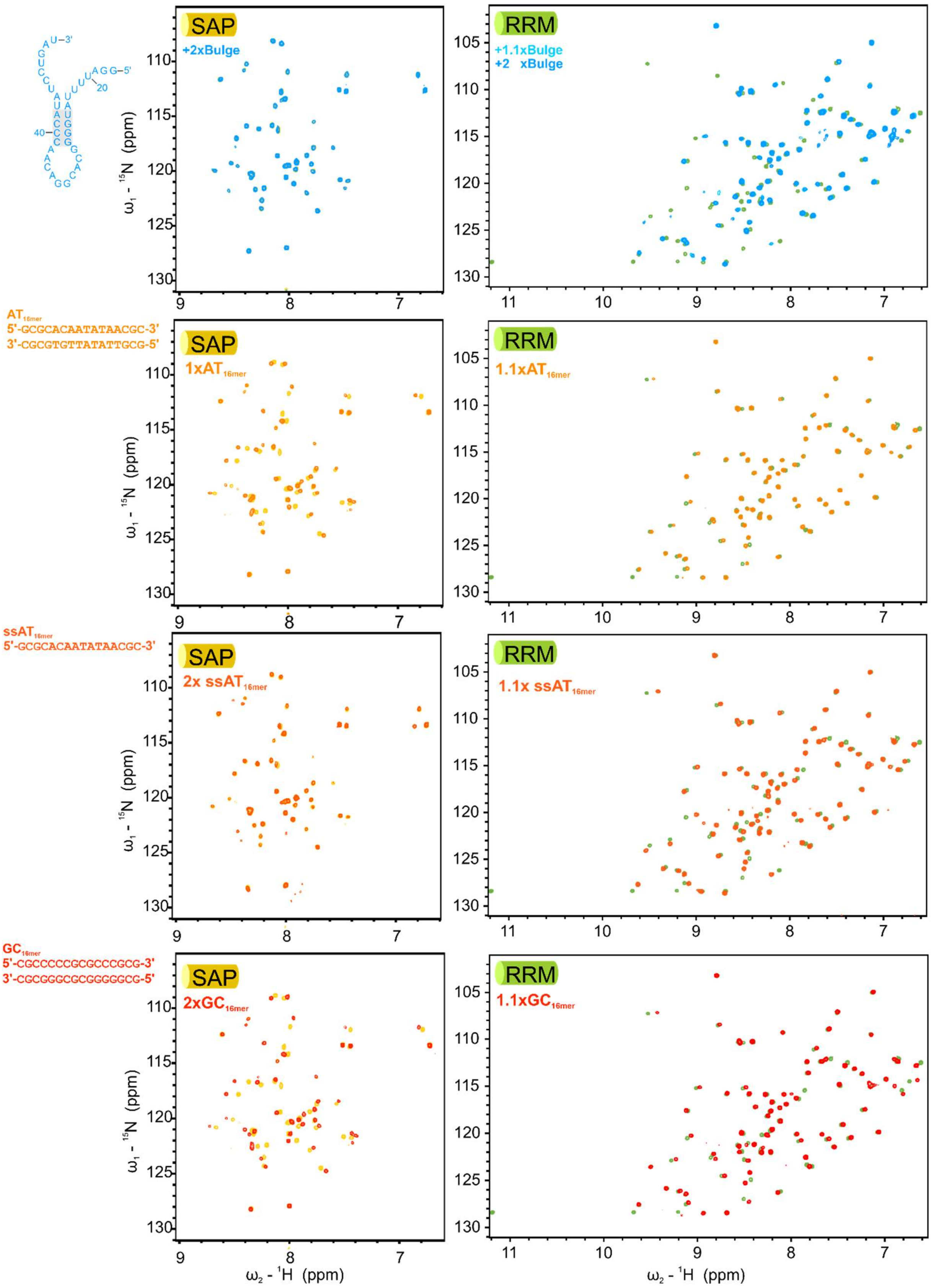

SAP
+2xBulge
RRM
+1.1xBulge
+2 xBulge
AT16mer
5'-GCGCACAATATAACGC-3'
3'-CGCGTGTTATATTGCG-5'
1xAT16mer
1.1xAT16mer
ssAT16mer
5'-GCGCACAATATAACGC-3'
2x ssAT16mer
1.1x ssAT16mer
GC16mer
5'-CGCCCCCGCGCCCCGCG-3'
3'-CGCGGGCGCGGGGGCG-5'
2xGC16mer
1.1xGC16mer
ω1 - 15N (ppm)
ω2 - 1H (ppm)

B

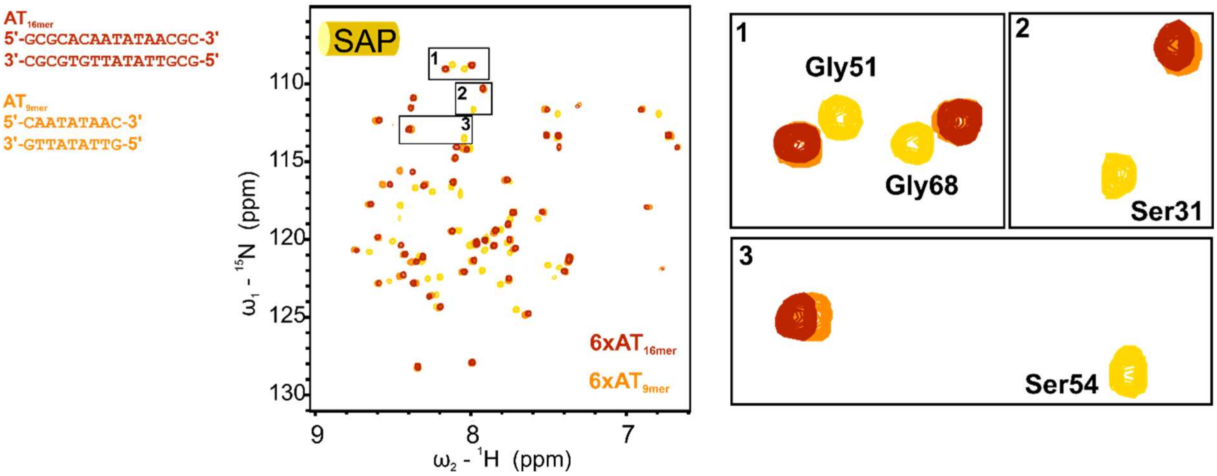


**Supplementary Figure 5:** Binding preferences of the SAFB2 SAP and RRM domains for different nucleic acid types. **(A)** Overlays of full $^{1}H$-$^{15}N$ HSQC spectra of the apo SAP domain (left column) or the RRM domain (right column) with spectra of the proteins in complex with respective molar excess (as indicated) of either Bulge RNA (purple, middle left), $dsAT_{16mer}$, $ssAT_{16mer}$ or $dsGC_{16mer}$ (from top to bottom). **(B)** Overlay of the full $^{1}H$-$^{15}N$ HSQC apo SAP spectrum with those of SAP in presence of molar excess of $dsAT_{16mer}$ (red) and $dsAT_{9mer}$ (orange). Zoom-ins of HSQC regions 1, 2 and 3 (right) reveal identical CSP trajectories for the 9mer and the 16mer dsDNA indicating the same binding mode.

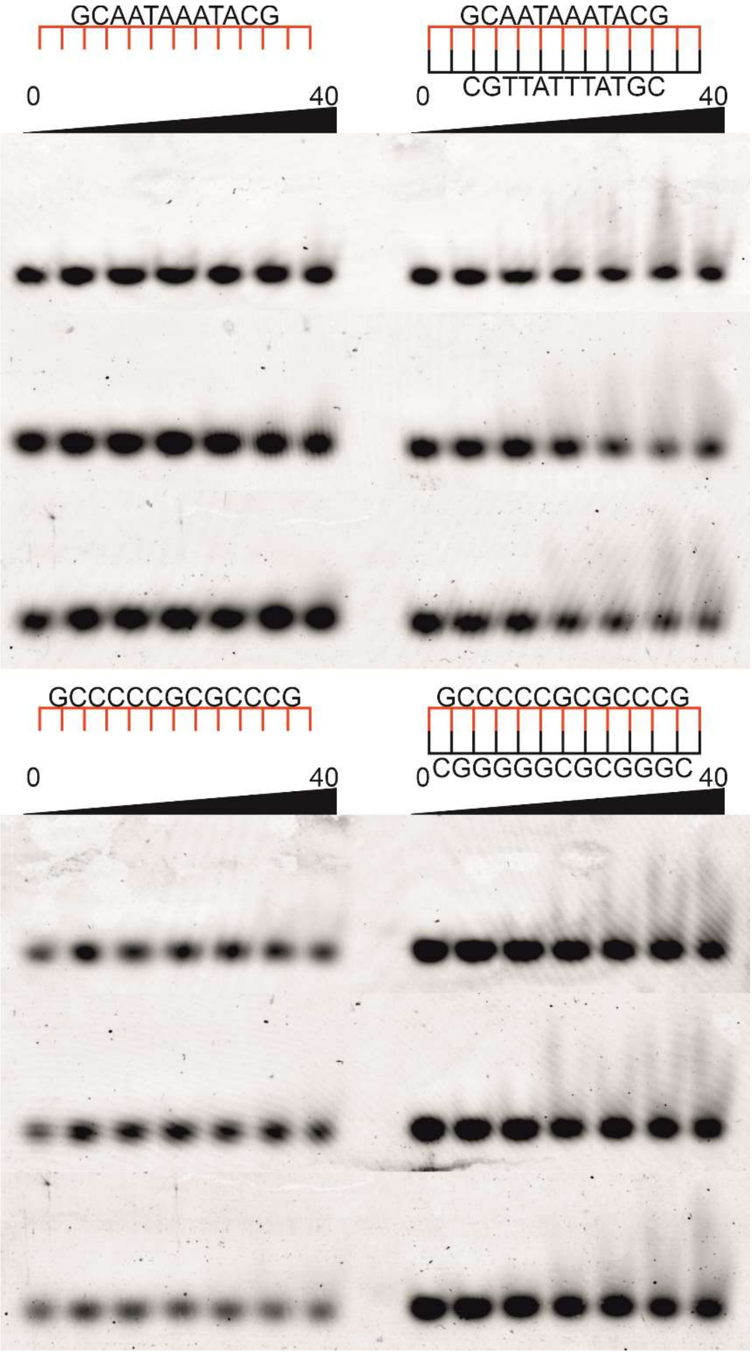


**Supplementary Figure 6:** Depiction of EMSA triplicates of the SAFB2 SAP domain titrated to single-stranded (left) and double-stranded (right) 13mer AT-rich or GC-rich DNAs (top and bottom, respectively) corresponding to main text Figure 5. Increasing protein concentrations (0-40 μM) are indicated by black bars above gel pictures.

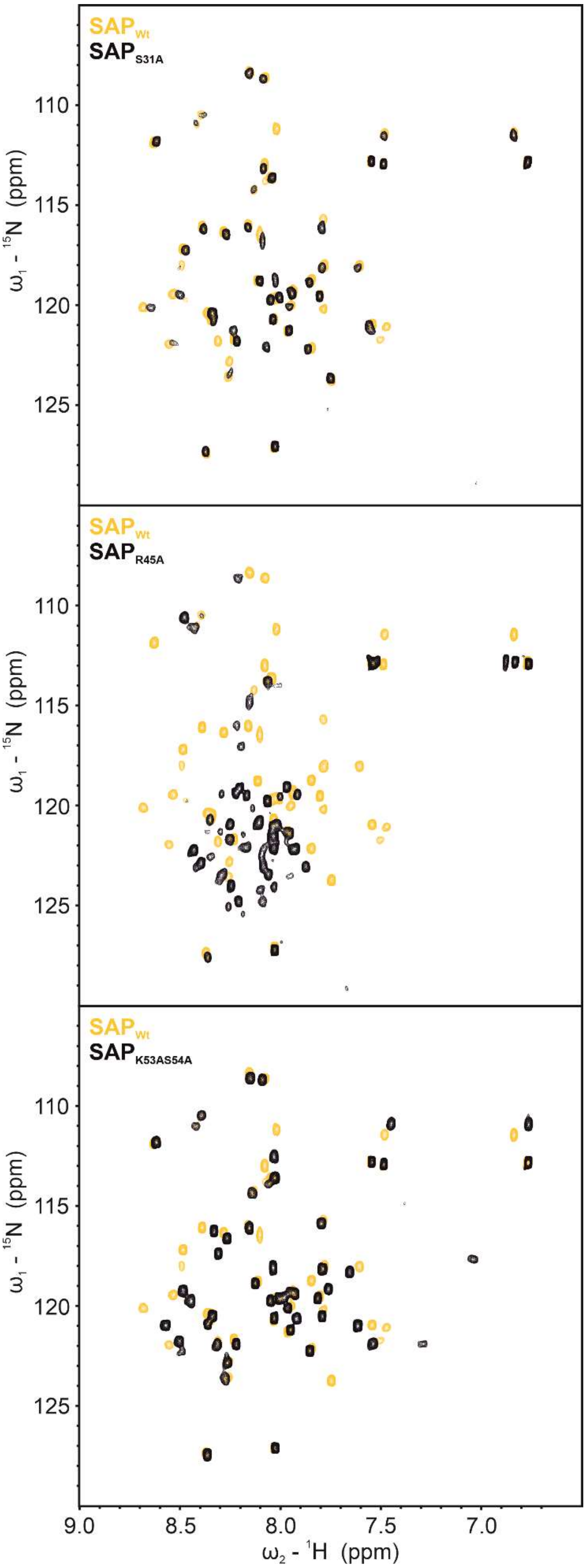


**Supplementary Figure 7:** Effects of individual point mutations on the SAFB2 SAP domain fold. Full $^{1}H$-$^{15}N$ HSQC spectra of mutant proteins: $SAP_{S31A}$, $SAP_{R45A}$ and $SAP_{K53AS54A}$ (from top to bottom), each overlaid with $SAP_{WT}$ for comparison.

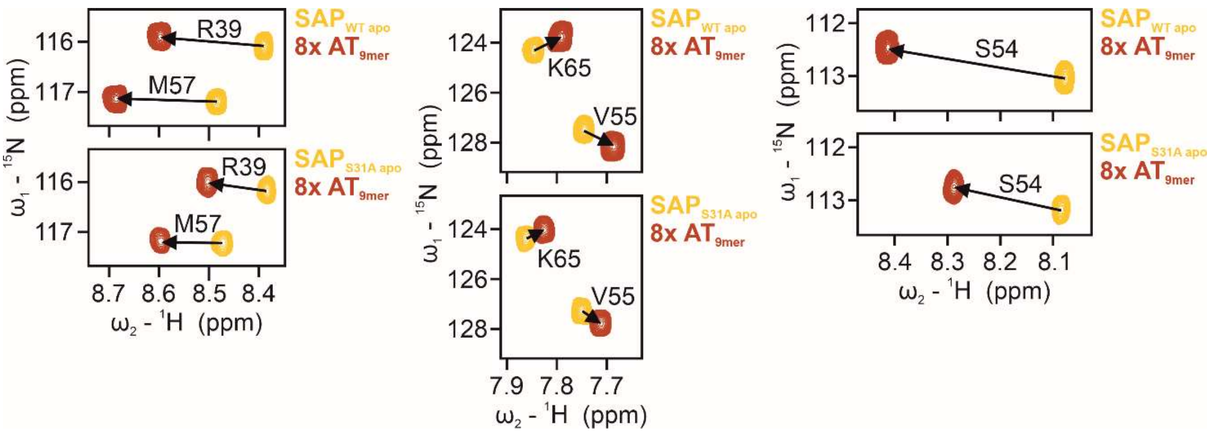


**Supplementary Figure 8:** Effect of the S31A mutation in the SAFB2 SAP domain on DNA binding. Identical insets of $^{1}H$-$^{15}N$ HSQC spectra of the $SAP_{WT}$ (upper panels) and the mutant $SAP_{S31A}$ (lower panels). Each overlay shows the apo protein (yellow) vs. complexes with 8- fold excess of $AT_{9mer}$ (red). Arrows indicate their respective CSP trajectories (for the selected residues). While trajectories are the same, the S31A mutant displays significantly reduced CSP extents. Consequently, the mutation reduces the affinity of SAP for the DNAs, but does not alter the mode of interaction or prohibit complex formation.

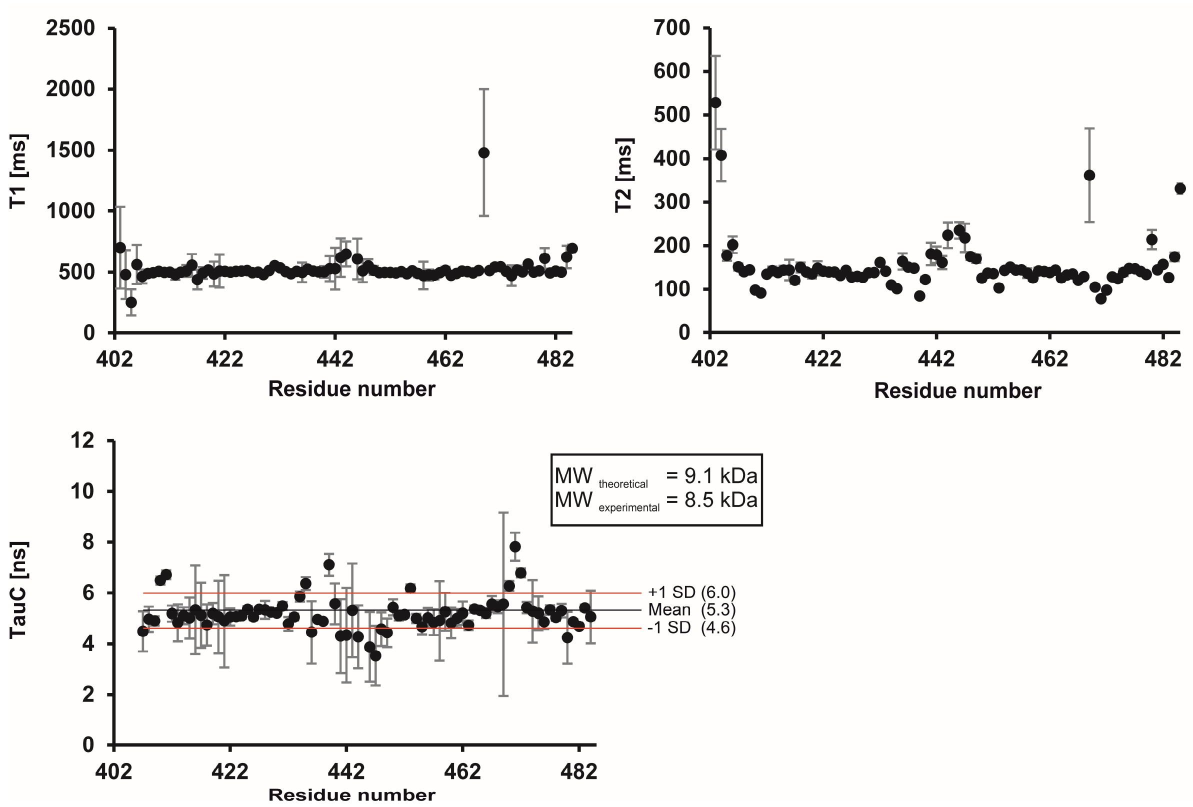


**Supplementary Figure 9:** $^{15}$N relaxation data for amides of the SAFB2 RRM domain. The upper two panels show $^{15}$N T1 and T2 times as a function of SAFB2 RRM domain amides recorded at an Avance II Bruker spectrometer (600 MHz proton Larmor frequency) and 298 K. For T1 and T2 all values are shown. Rotational correlation times TauC of the lower panel are derived from T1 and T2, but were calculated only for rigid parts according to hetNOE and SCS (see main text Figure 4) values and are shown with mean value ± 1 SD. T1 and T2 errors are determined by the program *CCPNMR Analysis;* TauC errors are the sum of percental errors obtained for T1 and T2.

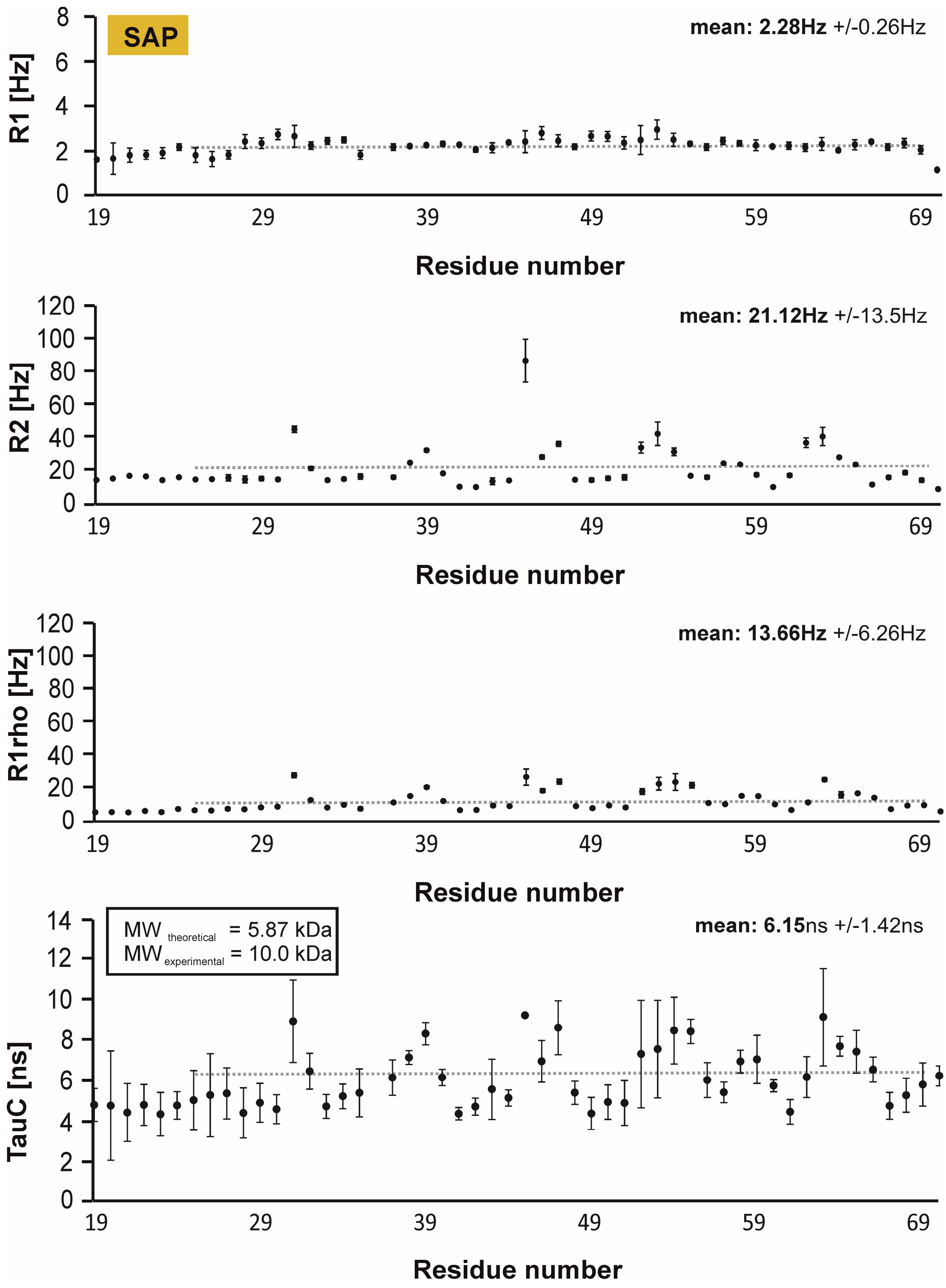


**Supplementary Figure 10:** $^{15}N$ relaxation data for the SAFB2 $SAP_{WT}$. R1, R2 and R1rho rates were determined from measurements at 700 MHz (proton Larmor frequency) and 298 K. Mean values were calculated for the structured domain indicated by grey, dotted lines, respectively. Errors for relaxation rates are derived from calculations using *CCPNMR Analysis* [9]. Mean TauC values (rotational correlation times) were used to estimate the SAP domain molecular weight ($MW_{experimental}$) and compared to the theoretical $MW_{theoretical}$ (as given in the box). Errors for TauC are the sum of percental errors obtained for R1 and R1rho.

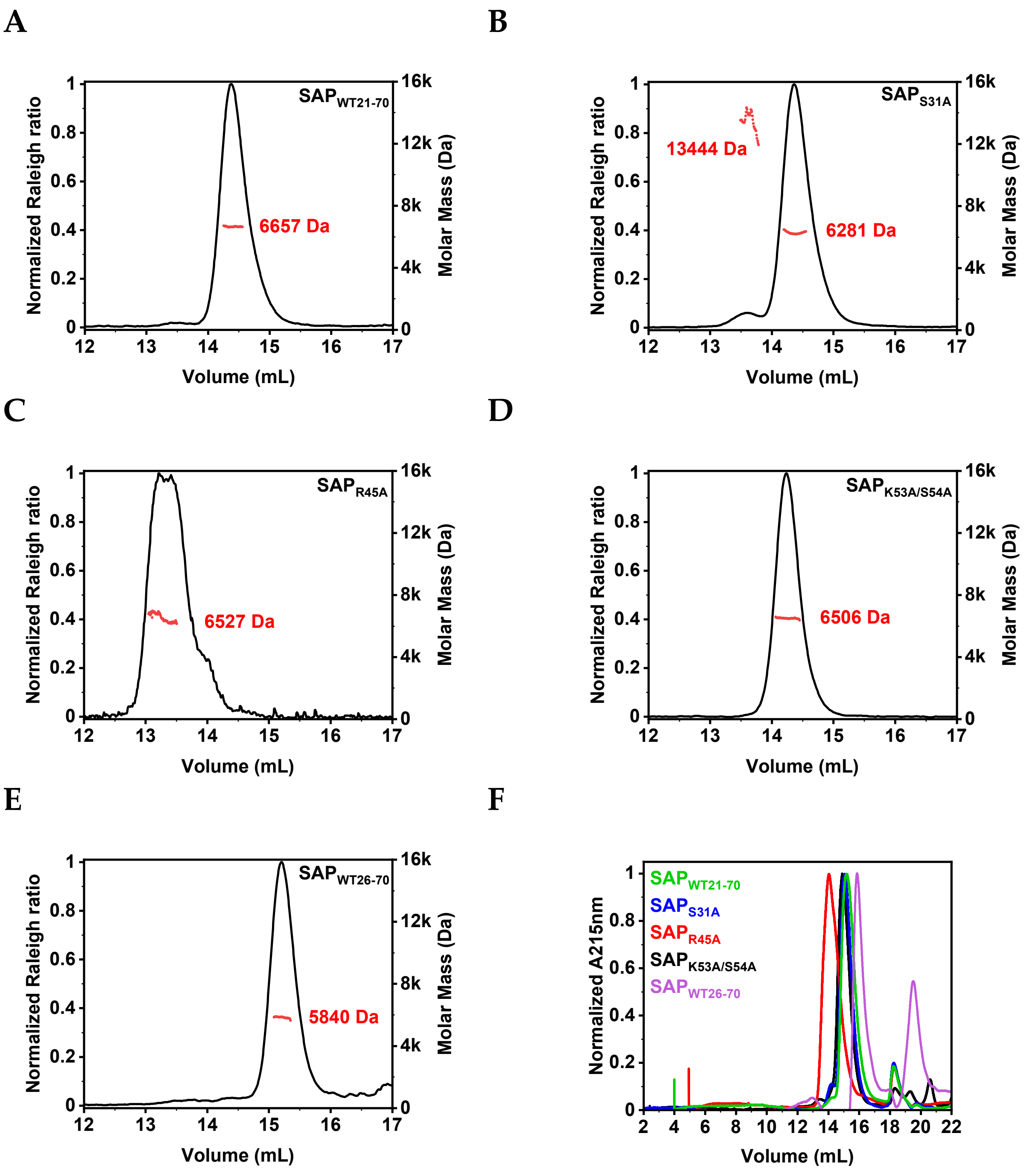


G

| | SAP$_{WT21-70}$ | SAP$_{31A}$ | SAP$_{R45A}$ | SAP$_{K53A/S54A}$ | SAP$_{WT26-70}$ |
|---|---|---|---|---|---|
| Theoretical MW (Da) | 5869 | 5853 | 5784 | 5796 | 5152 |
| aSEC based MW (Da) | 9810 | 10360 | 14857 | 10801 | 7620 |

**Supplementary Figure 11:** Analytical SEC and SEC-MALS data on SAP wild-type and mutants. **(A-E)** SEC-MALS analysis of SAP$_{WT21-70}$, SAP$_{31A}$, SAP$_{R45A}$, SAP$_{K53A/S54A}$ and SAP$_{WT26-70}$, respectively. Shown are the SEC chromatograms with the peak of interest and the MW as derived from the intersected peak along the elution volume as indicated by the red trace. **(F)** Analytical SEC of SAP wild-type and mutants. Full chromatograms corresponding to main text Figure 7. **(G)** Molecular weight estimation of SAP wild-type and mutants derived from a calibration curve based on elution volumes of MW standards when compared to the theoretical MWs of the respective monomers. All SEC runs in this figure were carried out on a SD75 Increase 10/300 GL column.

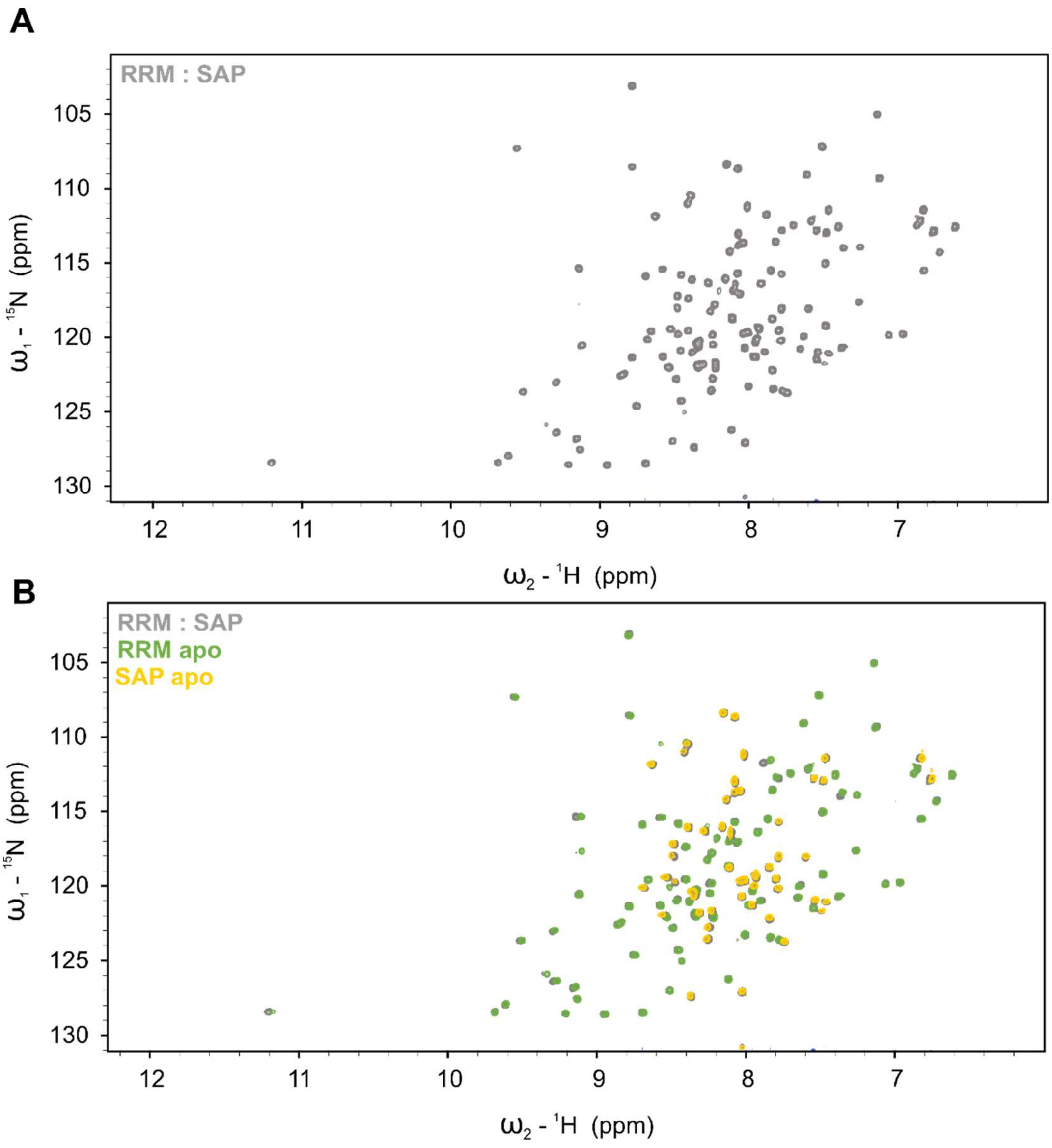


**Supplementary Figure 12:** The SAFB2 SAP and RRM domains do not interact. **(A)** $^{1}H$-$^{15}N$ HSQC of a mixed sample of $^{15}N$-labelled SAP and $^{15}N$-labelled RRM domains in a 1:1 ratio. **(B)** Overlay of the $^{1}H$-$^{15}N$ HSQC spectra of the 1:1 SAP/RRM mix from panel A with the respective spectra of individual $^{15}N$-labelled domains (RRM – green; SAP – yellow). The unperturbed peak patterns of either domain in the mixture indicate the two domains to be independent from each other.
